# GeneMAP: A discovery platform for metabolic gene function

**DOI:** 10.1101/2023.12.07.570588

**Authors:** Artem Khan, Gokhan Unlu, Phillip Lin, Yuyang Liu, Ece Kilic, Timothy C. Kenny, Kıvanç Birsoy, Eric R. Gamazon

## Abstract

Organisms maintain metabolic homeostasis through the combined functions of small molecule transporters and enzymes. While many of the metabolic components have been well-established, a substantial number remains without identified physiological substrates. To bridge this gap, we have leveraged large-scale plasma metabolome genome-wide association studies (GWAS) to develop a multiomic Gene-Metabolite Associations Prediction (GeneMAP) discovery platform. GeneMAP can generate accurate predictions, even pinpointing genes that are distant from the variants implicated by GWAS. In particular, our work identified *SLC25A48* as a genetic determinant of plasma choline levels. Mechanistically, SLC25A48 loss strongly impairs mitochondrial choline import and synthesis of its downstream metabolite, betaine. Rare variant testing and polygenic risk score analyses have elucidated choline-relevant phenomic consequences of SLC25A48 dysfunction. Altogether, our study proposes SLC25A48 as a mitochondrial choline transporter and provides a discovery platform for metabolic gene function.

## Main

Metabolic reactions are central to life, playing critical roles in energy production, nutrient absorption, waste removal and biomass synthesis. Given these critical processes, approximately 20% of protein-coding genes are dedicated to maintaining the intracellular chemical landscape and include small molecule transporters and enzymes. While decades of research have revealed the functions of a substantial number of these genes, the exact molecular substrates for many metabolic components remain elusive^1–8^. Such gaps in our understanding arise partly from the diverse tissue-specific expression patterns, functional redundancies and metabolic promiscuity of these elements, complicating efforts to define their precise physiological roles^9^. Dysfunction in metabolic functions is commonly associated with a range of disorders, from congenital anomalies to neurodegeneration and cancer^10–12^. Recent large-scale Genome-Wide Association Studies (GWAS) of the human metabolome have revealed pervasive genetic influences underlying chemical individuality^13–16^. Although these studies mostly focus on the mediating effects of metabolites on health outcomes^16–18^, such datasets also provide an opportunity to understand gene function. Yet, the main methodological challenge for using GWAS has been linking the phenotype-associated genetic variants to the relevant gene^19–21^.

To address this, we developed Gene-Metabolite Association Prediction (GeneMAP) Platform for discovery of metabolic gene function that leverages genetic models of gene expression and quantifies the gene-mediated genetic control of metabolites. To identify gene-metabolite relationships, we conducted Transcriptome-Wide Association Studies (TWAS)^22,23^ in two independent genomic studies of the human metabolome from CLSA^15^ and METSIM^14^ (Fig. 1a, Methods). The analysis yielded 526,934,749 gene-metabolite entries (26,956,587 unique gene-metabolite pairs) across all expression models that were matching between the two datasets (Fig. 1b). We then computed the *π_1_* statistic for each expression tissue model separately and assessed replicability of our findings^24,25^. Remarkably, the identified gene-metabolite associations were highly replicable with minimum *π_1_* > 0.8 (Fig. 1c). In agreement with the larger sample size for CLSA (n=8,299)^15^, *π_1_* was greater when CLSA was used as the discovery and METSIM as the validation dataset (Extended Fig. 1a). Using CLSA as the discovery dataset, we identified 102,058 significant gene-metabolite associations (q-value < 0.05) across all expression models, consisting of 12,041 unique pairs (Fig. 1d). Of note, the replicability weakly correlated with the GTEx tissue sample size^23^ used for the model training (Extended Fig. 1b). In contrast, the number of significant associations was well-correlated with tissue sample size^23^, demonstrating the importance of collecting large-scale datasets for enhanced detection power (Extended Fig. 1c). Interestingly, a proportion of gene-metabolite pairs were identified as significant only in a small number of tissue models, highlighting the utility of conducting multi-tissue model analysis (Extended Fig. 1d). The metabolite class categorization for the significant associations was consistent with their representations among the profiled compounds (Fig. 1e)^14^. Similar to CLSA single-SNP analysis where genomic loci displayed a low degree of pleiotropy, most of the genes were associated with few metabolites (Extended Fig. 1e)^15^.

**Figure 1.**
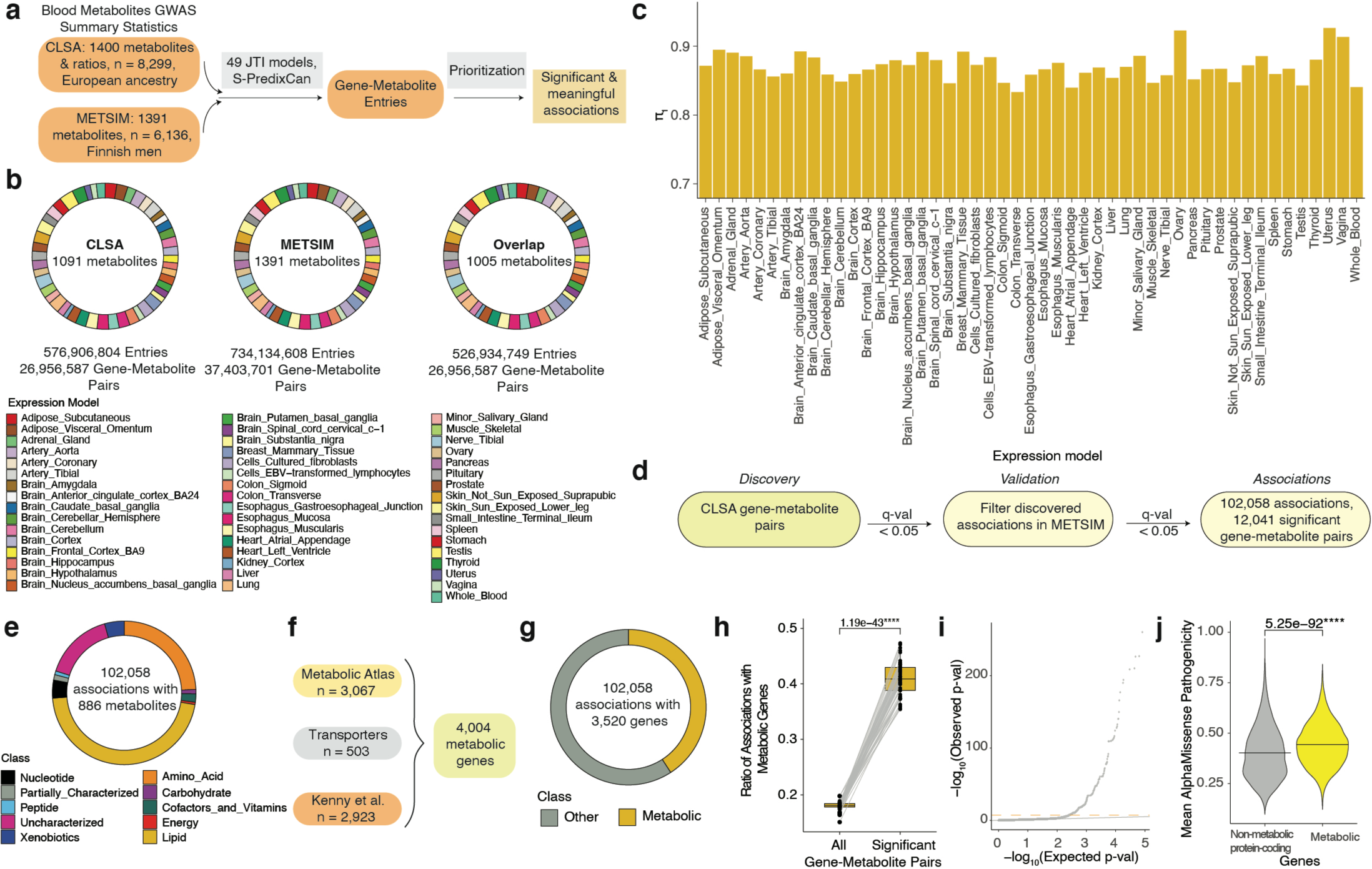
Summary and characterization of the gene-metabolite associations. a. Schematic for generating GeneMAP gene-metabolite associations. b. Summary of the generated gene-metabolite entries across 49 JTI expression models from the blood metabolite GWAS summary statistics in CLSA, METSIM, and the common gene-metabolite entries between the two databases (overlap). c. Bar graph displaying the replicability measure *π*_1_ across JTI expression models when CLSA was used as the discovery dataset and METSIM as the validation dataset. d. Pipeline for identification of significant gene-metabolite associations. e. Pie chart showing categorization of the gene-metabolite associations based on the metabolite class. f. Compiled list of metabolic genes based on the Metabolic atlas, study by Kenny et al., and set of transporters. g. Pie chart showing classification of the gene-metabolite associations based on gene subset (metabolic or other). h. Boxplot showing proportion of significant gene-metabolite pairs among all and metabolic genes across JTI expression models. Individual datapoints represent each JTI expression model. Lines connect the corresponding JTI expression model. Statistical significance was determined by two-tailed paired *t*-test. i. Representative QQ plot (whole blood JTI model) displaying the observed and expected distribution of −log_10_(METSIM p-value) for the gene-metabolite entries with metabolic genes that pass threshold p < 0.05 in CLSA. Expected distribution was calculated based on all the considered entries. The grey line is y = x, the orange dashed line is the Bonferroni-threshold based on gene-metabolite associations with all genes. j. Violin plot showing the distribution of the average AlphaMissense pathogenicity score for the genes in the non-metabolic protein coding (gray) and metabolic protein-coding genes (yellow) groups. Statistical significance was determined by two-tailed unpaired *t*-test.

We next asked whether the subset of metabolic genes could be a strong determinant of blood chemical composition (Fig. 1f). Indeed, we observed that the metabolic genes were enriched among the significant associations compared to all entries by two-fold in every tissue model set (Fig. 1g-h). This is also confirmed by the p-value distribution of the associations involving metabolic genes (Fig. 1i). Interestingly, AlphaMissense analysis showed that the average predicted pathogenicity of missense variants in the metabolic genes was significantly higher compared to non-metabolic genes (Fig. 1j)^26^. Consistent with this observation, CADD analysis revealed that the most deleterious variants in metabolic genes were more pathogenic compared to genome background and non-metabolic genes (Extended Fig. 1f-g)^27^. These findings further corroborate the critical role of metabolic genes in cellular and organismal processes. Altogether, the GeneMAP platform generated highly replicable and functionally informative results, with the potential to uncover novel biology.

Given the high replicability of the identified gene-metabolite associations, we next investigated biologically relevant complex structures such as Genetically Determined Metabolic Networks (GDMNs). For the nodes of these networks, we utilized 415 metabolites present among the significant associations with metabolic genes in whole blood in the discovery dataset. Our approach differs from conventional single-SNP GWAS methods through its focus on the genetically determined molecular traits. To build the networks, we computed metabolite-metabolite Pearson correlations and the corresponding empirical p-values from the selected gene-metabolite pairs’ effect sizes (β-values) (Fig. 2a). Notably, the correlations calculated from the CLSA and METSIM GeneMAP results were highly replicable (Fig. 2b). Furthermore, the pairs with empirical Bonferroni adjusted p-value < 0.05 computed from CLSA were enriched in the significant part of the p-value distribution determined from the METSIM dataset (Fig. 2c).

**Figure 2.**
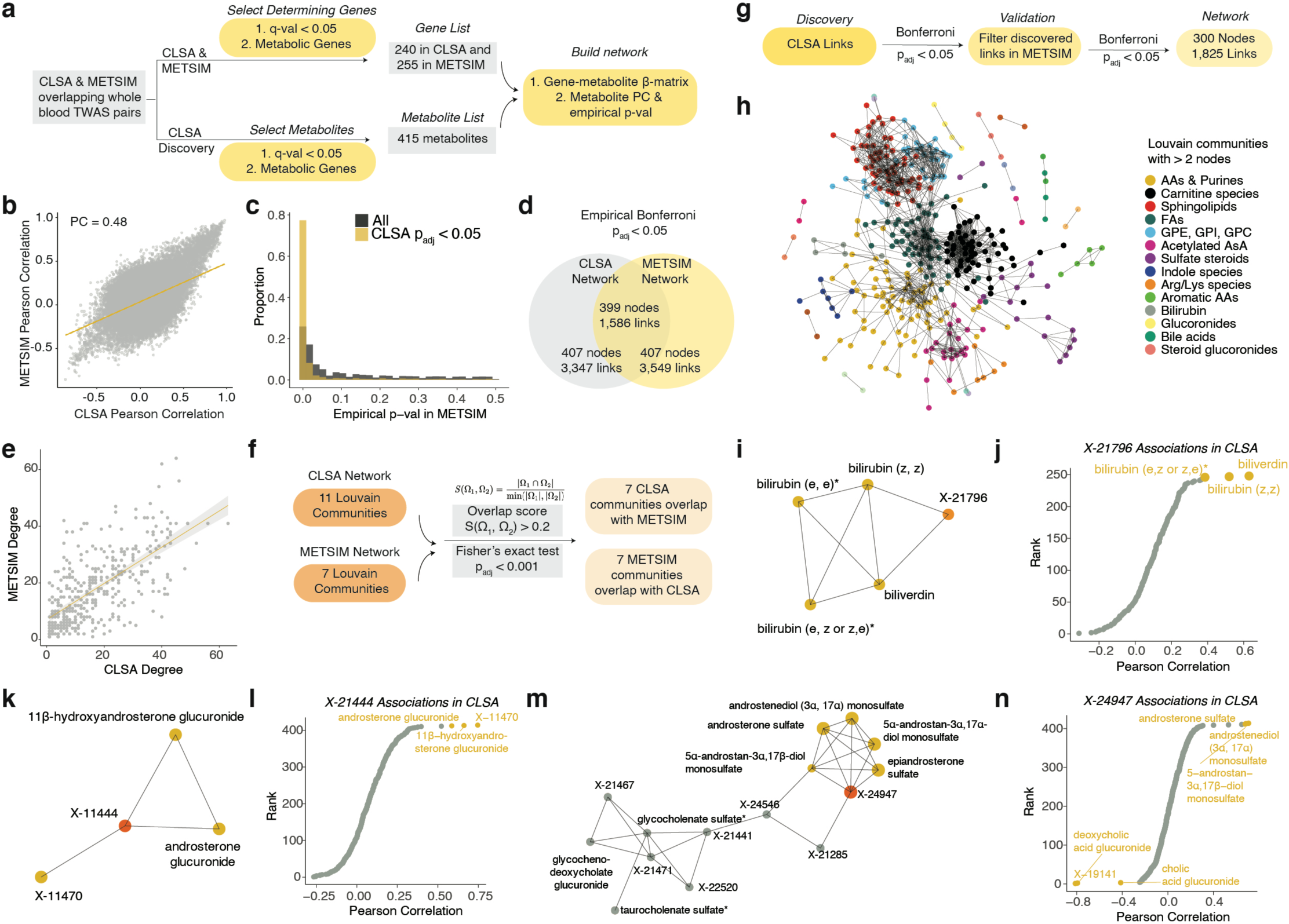
Genetically Determined Metabolic Networks (GDMNs) are replicable and interpretable. a. Schematic for the GDMNs construction pipeline for the whole blood JTI gene-metabolite associations. b. Comparison of Pearson correlations for the metabolite-metabolite pairs computed from the METSIM and CLSA datasets. c. Distribution of the empirical p-values in the validation (METSIM) dataset for all (gray) and only significant (Bonferroni adjusted p-value < 0.05, yellow) in discovery metabolite-metabolite pairs. d. Comparison of the links and edges in GDMNs built from the CLSA and METSIM datasets. e. Correlation of the node degrees in GDMNs built from the CLSA and METSIM datasets. f. Overlap between the Louvain communities in GDMNs from the CLSA and METSIM datasets. g. Schematic for building the consensus GDMN. h. Consensus GDMN with the Louvain community analysis. i. Louvain community annotated as Bile Acids from the consensus GDMN. j. Metabolite-metabolite pairs involving the uncharacterized compound X-21796. Data are plotted as ranks vs. Pearson correlations for the given metabolite-metabolite pair computed from the discovery dataset (CLSA). k. Louvain community annotated as glucoronides from the consensus GDMN. l. Metabolite-metabolite pairs involving the uncharacterized compound X-21444. Data are plotted as ranks vs. Pearson correlations for the given metabolite-metabolite pair computed from discovery (CLSA). m. Louvain community annotated as sulfate steroids from the consensus GDMN. n. Metabolite-metabolite pairs involving the uncharacterized compound X-24947. Data are plotted as ranks vs. Pearson correlations for the given metabolite-metabolite pair computed from discovery (CLSA).

To construct the edges of the GDMNs, we filtered the metabolite-metabolite entries using empirical Bonferroni corrected p-value < 0.05. GDMNs built from CLSA and METSIM resembled each other with almost all nodes and nearly half of the links being shared (Extended Fig. 2a-b, Fig. 2d). Strikingly, both local (e.g., node degrees) and global (e.g., Louvain communities) network properties were also conserved (Fig. 2e-f). To generate a consensus network from the two independent GDMNs, we filtered the edges based on the empirical p-values (Bonferroni adjusted p-value < 0.05) of the discovery and validation datasets. This consensus GDMN contained 300 nodes and 1,825 edges (Fig. 2g) with biochemically interpretable Louvain communities (Fig. 2h). As a biological network, the generated GDMN was scale-free based on the power-log distribution of node degree (Extended Fig. 2c)^28^. Moreover, it was resistant to random perturbations as indicated by a slower increase in the mean graph distance when an arbitrary node was removed as opposed to a hub (Extended Fig. 2d). Interestingly, this feature is in accordance with the properties of a previously reported metabolic network, which were speculated to underlie error tolerance to uniformly distributed mutations throughout evolution^28^. Remarkably, we could even infer the biochemical identity of some of the uncharacterized metabolites from the GDMN. For example, one of the five nodes of the bile acid Louvain community was an uncharacterized metabolite X-21796 suggesting a potential connection to bile acids (Fig. 2i). Consistent with this hypothesis, the top metabolite associations of X-21796 include bilirubin and biliverdin species implying that the metabolite is likely a bile acid derivative (Fig. 2j). Similarly, we predict X-21444 and X-24947 to be related to glucuronide and sulfate steroids species (Fig. 2k-n). Overall, the analysis of the complex structures of the GDMNs further demonstrates the high quality and replicability of the GeneMAP resource.

Given GeneMAP’s high replicability, we next asked whether our approach can both validate known gene-metabolite associations and discover novel ones. To enrich for metabolite determinants, we prioritized 2,245 (out of 12,041 entries) metabolic genes found to be significant in at least two expression models (Fig. 3a). Then, we tested whether these associations reflected known metabolite-associated loci as metQTLs from conventional single-SNP analysis and previously proposed effector genes. Notably, the relatively small subset of the prioritized GeneMAP findings covered 70% of the CLSA metQTLs (< 0.5 Mb) and 117 out of 145 effector genes (Fig. 3b-c)^15^. Interestingly, 564 of the 2,245 GeneMAP gene-metabolite pairs were distal (> 0.5 Mb) from the CLSA-reported metQTLs (Fig. 3d)^15^. This indicates that our approach not only recovers significant genetic signals mapped to effector genes but also identifies associations that single-SNP methods are relatively underpowered to detect.

**Figure 3.**
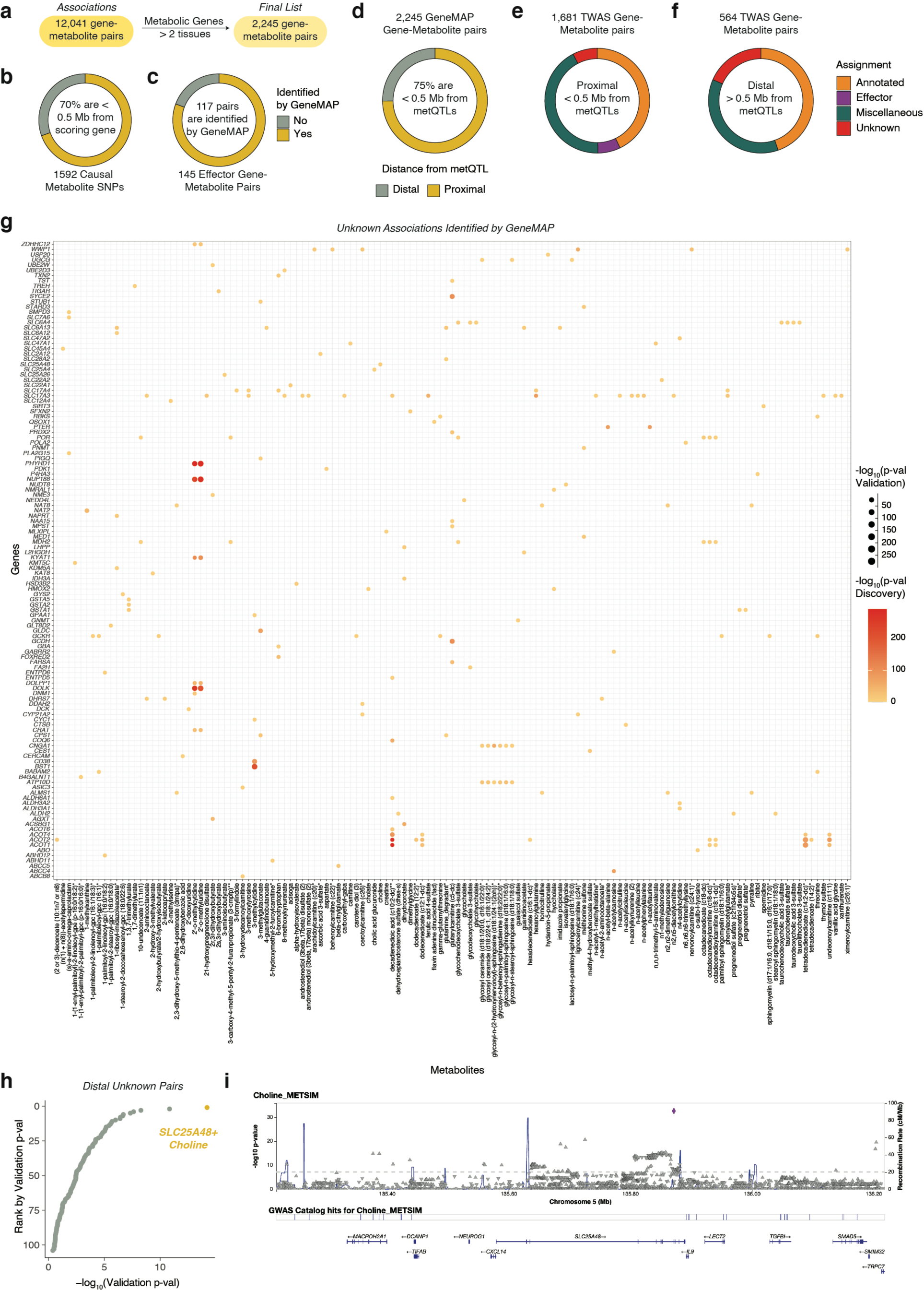
GeneMAP identifies *SLC25A48* as a genetic determinant of blood choline level. a. Schematic showing filtering and prioritization of the gene-metabolite associations. b. Pie chart displaying the proportion of the identified SNPs as causal in CLSA single-SNP analysis (Chen et al.) for the corresponding metabolite proximal (< 0.5 Mb) to the prioritized gene-metabolite GeneMAP associations. Only metabolites measured in both CLSA and METSIM were considered. Yellow represents identified by GeneMAP, gray – not. c. Pie chart displaying the proportion of the identified gene effector-metabolite pairs in CLSA single-SNP analysis (Chen et al.) by GeneMAP. Yellow represents identified by GeneMAP, gray – not. Only metabolites measured in both CLSA and METSIM were considered. d. Pie chart displaying the proportion of the 2,245 GeneMAP prioritized gene-metabolite proximal (yellow) or distal (gray) to identified SNPs for the corresponding metabolite in CLSA. e. Annotation of the GeneMAP gene-metabolite associations proximal (< 0.5 Mb) to the significant SNPs identified in CLSA single-SNP analysis. f. Annotation of the GeneMAP gene-metabolite associations distal (> 0.5 Mb) to the significant SNPs identified in CLSA single-SNP analysis. g. Bubble plot of unknown gene-metabolite associations identified by GeneMAP. Bubble color corresponds to the −log_10_(p-value) of the association between the indicated gene and metabolite in discovery (CLSA) dataset. Bubble size represents the −log_10_(p-value) of the association between the indicated gene and metabolite in validation (METSIM) dataset. h. GeneMAP undescribed associations distal from the SNPs identified in CLSA single-SNP analysis. Data are plotted as ranks vs. −log_10_(p-value) for the given gene-metabolite association in Validation (METSIM). i. LocusZoom plot for METSIM choline association signals in *SLC25A48* region.

To further assess the quality of the GeneMAP gene-metabolite pairs, we utilized literature curation and several genomics databases (Supplementary Table 1)^14–16,29,30^. Among 1,681 proximal associations, we could find supporting evidence for 720 (which we term “Annotated”) in addition to the 117 effector-metabolite entries identified in the single-SNP analysis of CLSA^15^ (Fig. 3e). For 128 pairs denoted as “Unknown” and potentially pointing to novel biology, no causal gene in the nearby genomic region could be nominated (Fig. 3e). The rest (“Miscellaneous”) included entries with uncharacterized and partially characterized metabolites, or with nearby nominated causal genes (Fig. 3e). Strikingly, we could provide supporting evidence for 252 out of 564 that were distal from the metQTL associations^15^ (Fig. 3f). This shows that GeneMAP captures current knowledge of gene-metabolite pairs, including those missed by a single-SNP analysis approach (“Undetected”).

To find novel associations, we focused on the list of “Unknown” entries (Fig. 3g) and sought to find functionally informative gene-metabolite pairs among the “Undetected” entries. We therefore ranked the GeneMAP associations distal from the previously reported (CLSA) metQTLs^15^ by the minimum validation (METSIM) p-value across all models. This analysis yielded the *SLC25A48*-choline pair as the top GeneMAP hit among the “Unknown” and “Undetected” associations (Fig. 3h). Whereas no variant within the *SLC25A48* genomic locus reached the genome-wide significance threshold in the CLSA dataset for choline, the METSIM dataset contained multiple significant findings (Fig. 3i, Extended Fig. 3a)^14^. Furthermore, the recent GCKD study provides additional support for *SLC25A48* variant associations with choline (Extended Fig. 3b)^13^. Using Mendelian randomization (MR-Egger regression and weighted median), we demonstrate the causal effect of *SLC25A48* on choline (p-value_MR-Egger_ < 10^-100^, p-value_median-based_ = 0.001, Extended Fig. 3c-d). Altogether, these findings are consistent with a greater detection power from GeneMAP for identifying novel gene-metabolite associations.

To test our prediction and uncover the causal links behind the gene-metabolite relationships, we focused on the top scoring association between *SLC25A48* and choline. Choline is involved in a wide range of physiological processes such as neurotransmission, methyl-group metabolism, and lipid metabolism^31^. In most cells, it has two major metabolic fates: in the cytosol, choline kinase alpha (CHKA) phosphorylates free choline to form phosphocholine, which is a precursor for the production of phosphatidylcholine, a major component of the cell membranes^32,33^. Additionally, free choline can enter mitochondria for a two-step oxidation to produce betaine, a key metabolite in one-carbon metabolism and an osmolyte^32,34–36^. Given that SLC25A48 is a member of SLC25A family that encompasses mitochondrial small molecule transporters, we hypothesized that SLC25A48 may regulate the availability of choline or its downstream metabolites in mitochondria.

To begin to study the function of SLC25A48 in cellular metabolism, we first asked whether loss of SLC25A48 leads to any change in the abundance of mitochondrial metabolites. We therefore generated HEK293T *SLC25A48*-knockout cells as well as those complemented with *SLC25A48* cDNA (Extended Fig. 4a-c). These cells also express 3xHA-OMP25 Mito-Tag, which enables immunopurification of mitochondria for metabolic profiling^37^. Consistent with the predicted cellular localization^38^, immunofluorescence experiments confirmed mitochondrial localization of SLC25A48 (Fig. 4a). Using these engineered cells, we next performed metabolomic profiling on immunopurified mitochondria by liquid chromatography-mass spectrometry (Fig. 4b, Extended Fig. 4d). While the abundance of most metabolites was similar, we observed a 4-fold drop of betaine and ∼50-fold depletion of choline in mitochondria from *SLC25A48*-knockout cells compared to cDNA-complemented controls (Fig. 4c). Notably, SLC25A48 loss did not alter the cellular phosphocholine availability, suggesting that it does not impact cytosolic choline metabolism or import (Extended Fig. 4e). To formally test whether SLC25A48 specifically is involved in production of betaine, but not phosphocholine, we conducted a whole cell [1,2-^13^C_2_]Choline isotope tracing experiment (Fig. 4d). Indeed, HEK293T lacking SLC25A48 had a significantly reduced incorporation of isotope-labeled choline into betaine, but not phosphocholine in two different single clones as well as compared to the parental cell line (Fig. 4e-f, Extended Fig. 4f-i). Altogether, these results suggest that SLC25A48 is a determinant of *de novo* betaine synthesis from choline.

**Figure 4.**
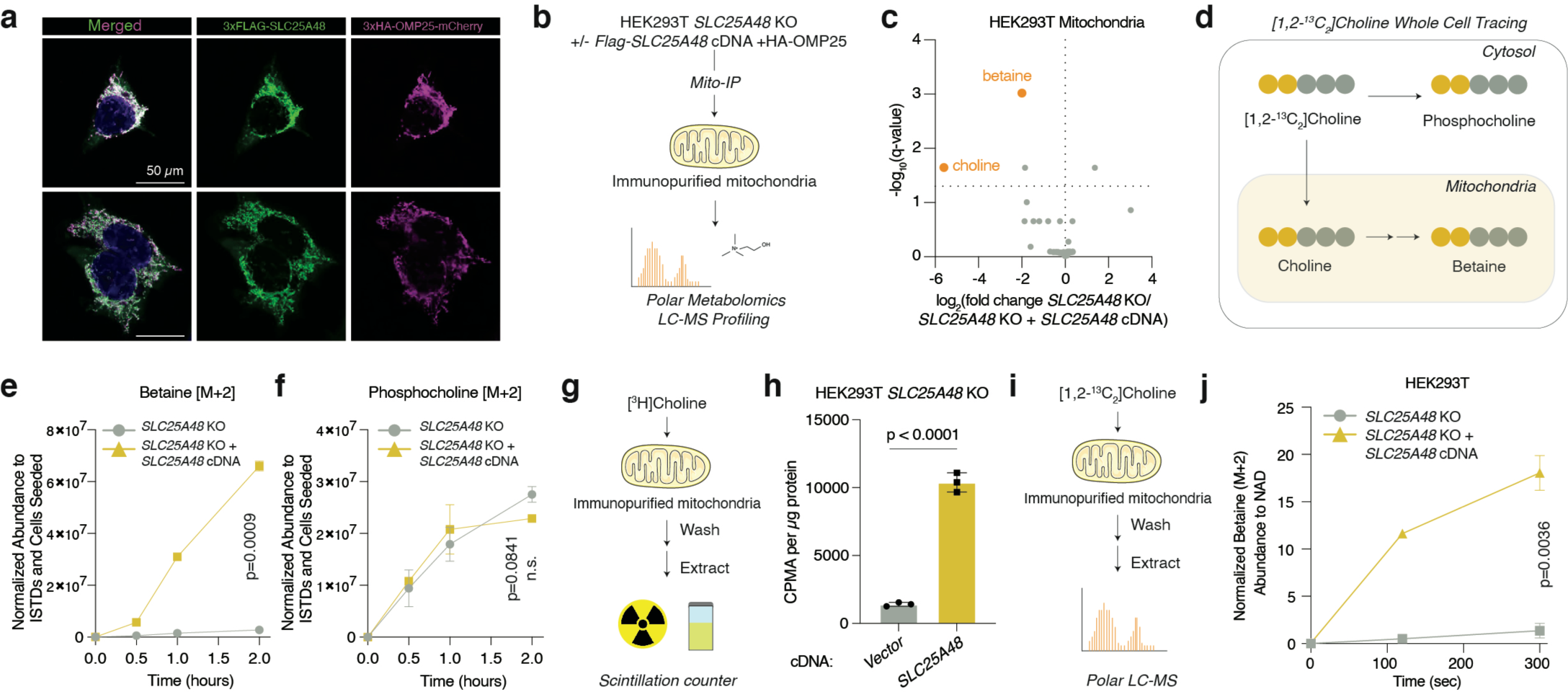
SLC25A48 mediates mitochondrial choline import. a. Immunofluorescence analysis of SLC25A48 (Flag, green), HA (HA, red) and nucleus (DAPI staining, blue) in HEK293T cells. Scale bar is 50 µm. b. Schematic for polar metabolomic profiling of immunopurified mitochondria from HEK293T *SLC25A48*-knockout cells expressing an empty vector control or *3xFLAG-SLC25A48* cDNA. c. Volcano plot with −log_10_(q-value) vs. log_2_ fold change in metabolite abundance normalized to isotope labeled amino acid standards (ISTDs) and NAD level in immunopurified mitochondria from HEK293T *SLC25A48*-knockout cells expressing a vector control or *SLC25A48* cDNA. The dotted line is the significance threshold of q-value < 0.05. Two-tailed unpaired t-tests followed by Benjamini, Krieger, and Yekutieli multiple test correction were performed. d. Schematic for the whole cell [1,2-^13^C_2_]Choline tracing. e. Betaine [M+2] abundance in HEK293T *SLC25A48*-knockout cells expressing an empty vector control or *3xFLAG-SLC25A48* cDNA after incubation with [1,2-^13^C_2_]Choline for the indicated time points. Data are mean ± standard deviation and normalized by ISTDs and cells seeded; n = 3. Statistical significance determined by two-way RM ANOVA followed by post hoc Bonferroni multiple correction. f. Phosphocholine [M+2] abundance in HEK293T *SLC25A48*-knockout cells expressing an empty vector control or *3xFLAG-SLC25A48* cDNA after incubation with [1,2-^13^C_2_]Choline for the indicated time points. Data are mean ± standard deviation and normalized by ISTDs and cells seeded; n = 3. Statistical significance determined by two-way RM ANOVA followed by post hoc Bonferroni multiple correction. g. Schematic for the 5-min [^3^H]Choline radioactive uptake in immunopurified mitochondria from HEK293T *SLC25A48*-knockout cells expressing an empty vector control or *3xFLAG-SLC25A48* cDNA. h. Uptake of [Methyl-^3^H]-Choline by immunopurified mitochondria from HEK293T *SLC25A48*-knockout cells expressing a vector control or *3xFLAG-SLC25A48* cDNA for 5 min. Data are mean ± standard deviation and normalized by ug of protein; n = 3. Statistical significance was determined by two-tailed unpaired t-test. i. Schematic for the [1,2-^13^C_2_]Choline tracing in immunopurified mitochondria from HEK293T *SLC25A48*-knockout cells expressing an empty vector control or *3xFLAG-SLC25A48* cDNA. j. Betaine [M+2] abundance in immunopurified mitochondria from HEK293T *SLC25A48*-knockout cells expressing an empty vector control or *3xFLAG-SLC25A48* cDNA incubated in [1,2-^13^C_2_]Choline uptake for the indicated timepoints. Data are mean ± standard deviation and normalized by NAD abundance; n = 3. Statistical significance determined by two-way RM ANOVA followed by post hoc Bonferroni multiple correction. All the experiments were repeated at least twice.

Betaine production involves the two-step oxidation of choline by two mitochondrially-localized enzymes (choline dehydrogenase and aldehyde hydrogenase 7 family member 1)^32,34–36^. Given this, we considered the possibility that SLC25A48 mediates mitochondrial choline import and thereby impacts betaine production. To determine this, we performed a radioactive [Methyl-^3^H]Choline uptake assay on immunopurified mitochondria from *SLC25A48*-knockout and cDNA-complemented controls (Fig. 4g). During a 5-minute incubation, mitochondria without SLC25A48 took up 7-fold less [Methyl-^3^H]Choline compared to the protein-containing control (Fig. 4h). To further confirm this, we conducted [1,2-^13^C_2_]Choline isotope tracing using immunopurified mitochondria (Fig. 4i). Indeed, we observed a substantial drop in the labeled choline incorporation into betaine in mitochondria from *SLC25A48*-knockout cells compared to cDNA-expressing controls (Fig. 4j). Altogether, our results suggest that *SLC25A48* is necessary for mitochondrial choline import and is a key determinant of *de novo* betaine synthesis in mammalian cells.

Given the cellular role of SLC25A48 in choline metabolism, we sought to identify the gene’s role in human health and disease. To evaluate the impact of SLC25A48 on the medical phenome, we conducted comprehensive rare variant analysis of 1,437 phenotypes in 469,787 individuals (UK Biobank), using the *SLC25A48* predicted loss-of-function (pLoF) variants (Fig. 5a)^39^. Restricting to such variants that completely disable the gene may enable new insights into pathophysiological processes. The departure of the observed p-value distribution from the expected (theoretical null) one identified highly significant disease associations (Fig. 5b), including musculoskeletal diseases and digestive disorders. Next, we examined whether the top disease phenotypes from the loss-of-function (LoF) analysis (p-value < 0.05) were also genetically determined by blood choline level. We tested the polygenic risk score (PRS) trained in the CLSA choline GWAS and found 17 significant associations with choline level among the top LoF implicated diseases (Fig. 5c-d). Among these, three had passed the stringent threshold of FDR < 0.05 from the LoF analysis: Ichthyosis congenita, hereditary disturbances in tooth structure, and diverticulosis (Fig. 5e). Overall, the combination of rare variant LoF testing and the choline PRS analysis identifies the potentially causal effects of *SLC25A48* in human disease via the mediated effects of choline.

**Figure 5.**
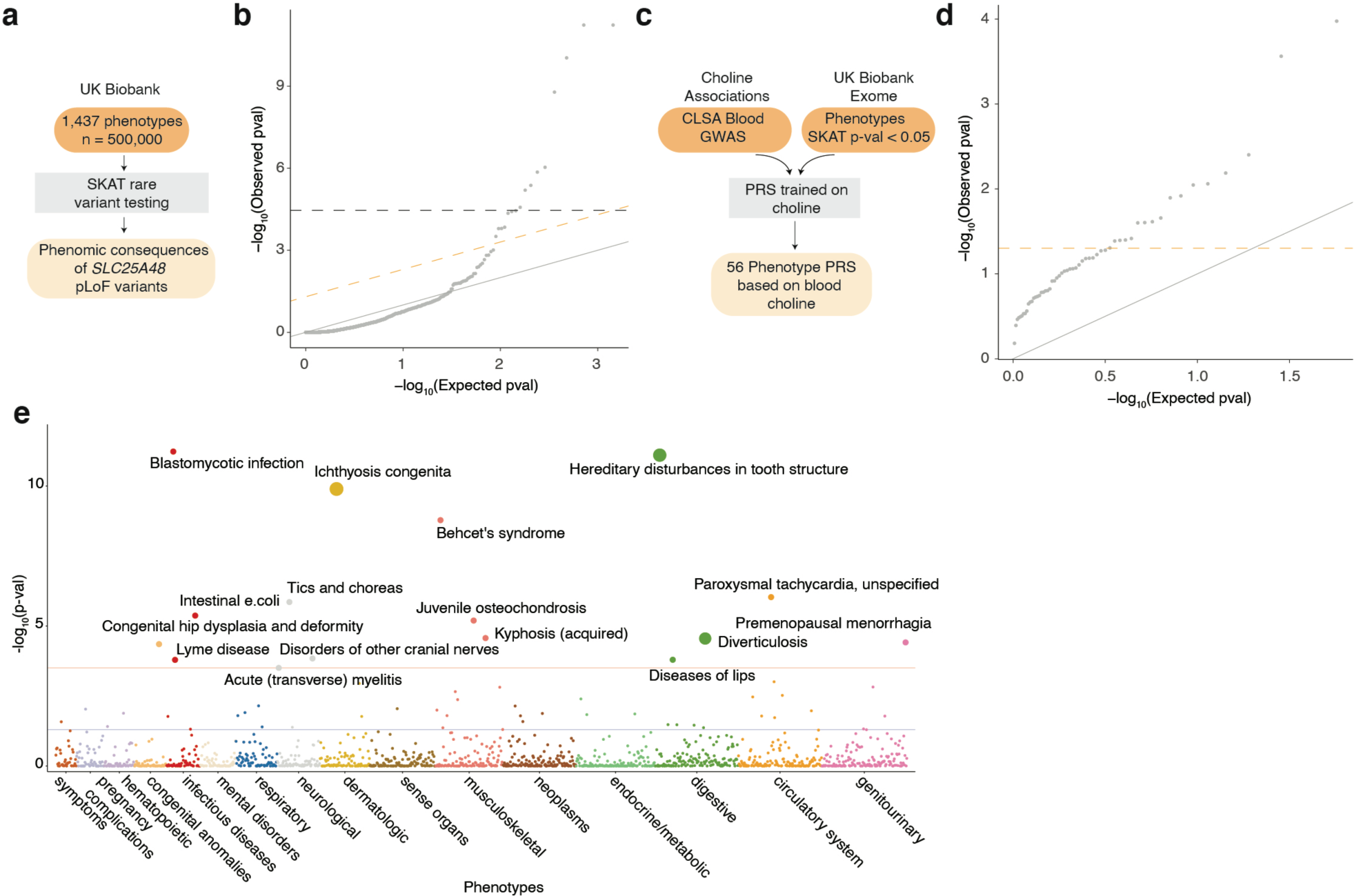
Choline-related phenomic consequences of SLC25A48 dysfunction. a. Pipeline for rare variant testing analysis on *SLC25A48* pLoF variants. b. QQ plot showing the observed and expected distribution of −log_10_(p-value) for the rare variant testing results on *SLC25A48* pLoF variants. The grey line is y = x, the orange dashed line corresponds to FDR = 0.05, the black dashed line is Bonferroni-threshold based on the number of considered phenotypes. c. Pipeline for phenotype polygenic risk score (PRS) calculations from choline GWAS. d. QQ plot showing the observed and expected distribution of −log_10_(p-value) for the PRS results on the significant phenotypes from the rare variant testing (p-value < 0.05). The grey line is y = x, the orange dashed line is the p-value = 0.05. e. Manhattan plot displaying phenotypes grouped by categories (x-axis) and p-value from the rare variant testing analysis on *SLC25A48* pLoF variants. The blue line is showing the threshold for p-value = 0.05, the red line for FDR = 0.05. Large size dots indicate phenotypes with rare variant testing FDR < 0.05 and PRS p-value < 0.05

In this study, we developed GeneMAP, a platform for predicting metabolic gene function, and demonstrated its ability to render accurate and replicable results. GeneMAP uncovers functionally informative gene-metabolite pairs distal from the GWAS implicated variants, which single-SNP analysis approaches are relatively underpowered to detect. In line with the robustness of our approach, we identified an orphan mitochondrial protein SLC25A48 as a mediator of choline mitochondrial import. Our data suggest that SLC25A48 is a major determinant of *de novo* synthesis of betaine, a critical metabolite for one-carbon metabolism and osmolyte produced from choline^32,34–36^. Furthermore, combined rare variant testing and polygenic risk score analysis revealed the novel choline-associated consequences of SLC25A48 dysfunction in health that include digestive and musculoskeletal disorders. Further work is necessary to elucidate the structural mechanisms underlying the precise role of SLC25A48 in mitochondrial choline import and its physiological relevance. Given that 30% of the SLC transporter family still do not have identified physiological substrates, GeneMAP provides a unique platform for deorphanizing many members of this family. Altogether, our work highlights the utility of a genetics-anchored multiomic approach for studying the role of a large class of genes in cellular and organismal metabolism. All generated results are publicly available through an interactive online portal for the use of the scientific community.

**Extended Figure 1.**
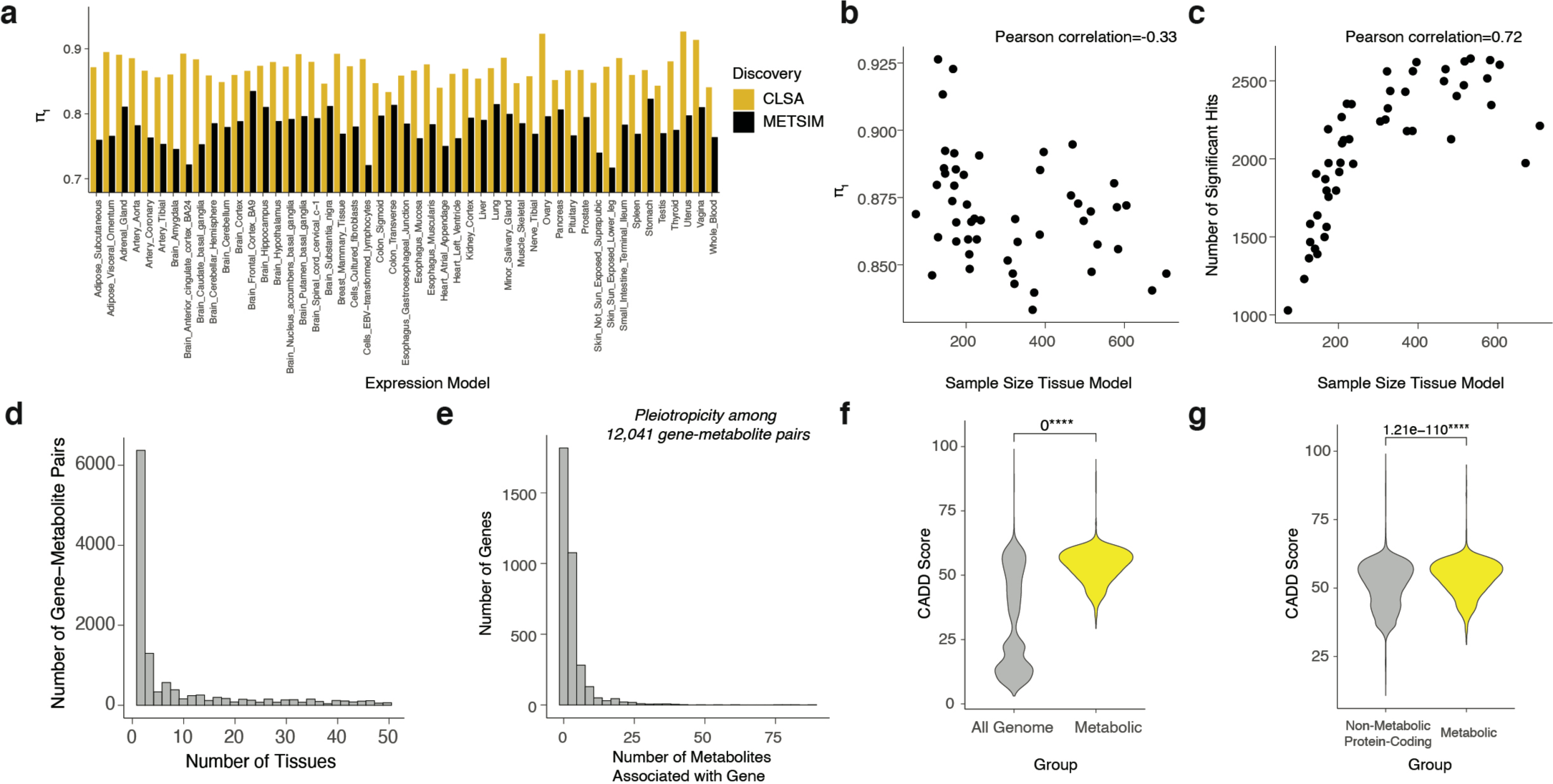
Development of the GeneMAP pipeline. a. Bar graph displaying the replicability measure *π*_1_ across JTI expression models when CLSA was used as discovery and METSIM as validation (yellow) and vice versa (black). b. Replicability measure *π*_1_ as a function of the tissue sample size in GTEx used for the JTI expression model training. c. Number of identified significant gene-metabolite associations as a function of the tissue sample size in GTEx used for the JTI expression model training. d. Histogram with the distribution of the number of JTI expression models in which the significant gene-metabolite associations are present. e. Pleiotropicity of the genes. Histograms with the distribution of the number of metabolites associated with the gene among the significant gene-metabolite associations. f. Violin plot showing the distribution of the top CADD scores for individual genes in the whole-genome (gray) and metabolic genes (yellow) groups. Statistical significance was determined by two-tailed unpaired *t*-test. g. Violin plot showing the distribution of the top CADD scores for the genes in the non-metabolic protein coding (gray) and metabolic genes (yellow) groups. Statistical significance was determined by two-tailed unpaired *t*-test.

**Extended Figure 2.**
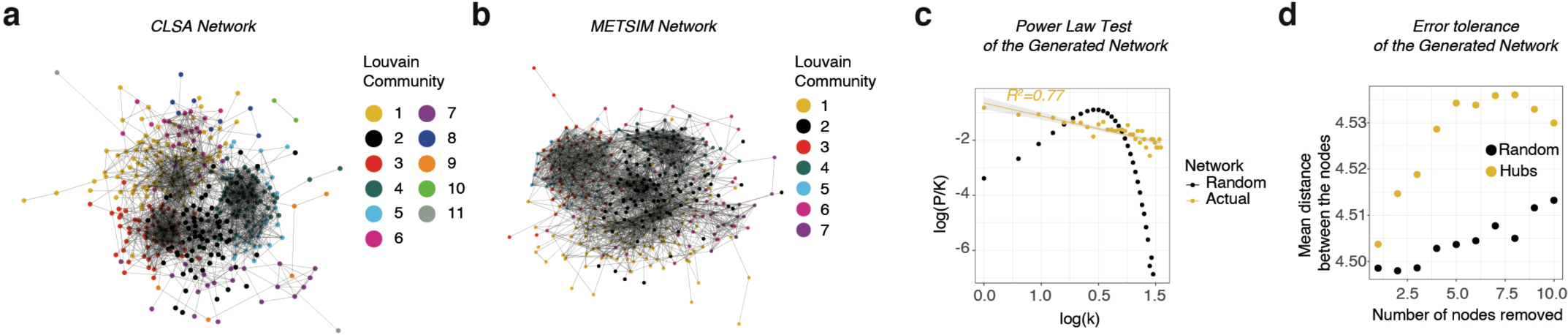
Genetically Determined Metabolic Network Analysis. a. GDMN built from CLSA dataset with the Louvain community analysis. b. GDMN built from METSIM dataset with the Louvain community analysis. c. Distribution of the node degrees for the observed consensus network (yellow points) and simulated random networks (n = 20,000, black points). The yellow line is regression line for the consensus network degree distribution with R^2^. d. Error-tolerance of the consensus GDMN represented as an average distance between the metabolic nodes in network when consequently removing 10 random nodes (black points, simulated 500 times) and 10 hubs (yellow points, ordered from the most to least connected nodes).

**Extended Figure 3.**
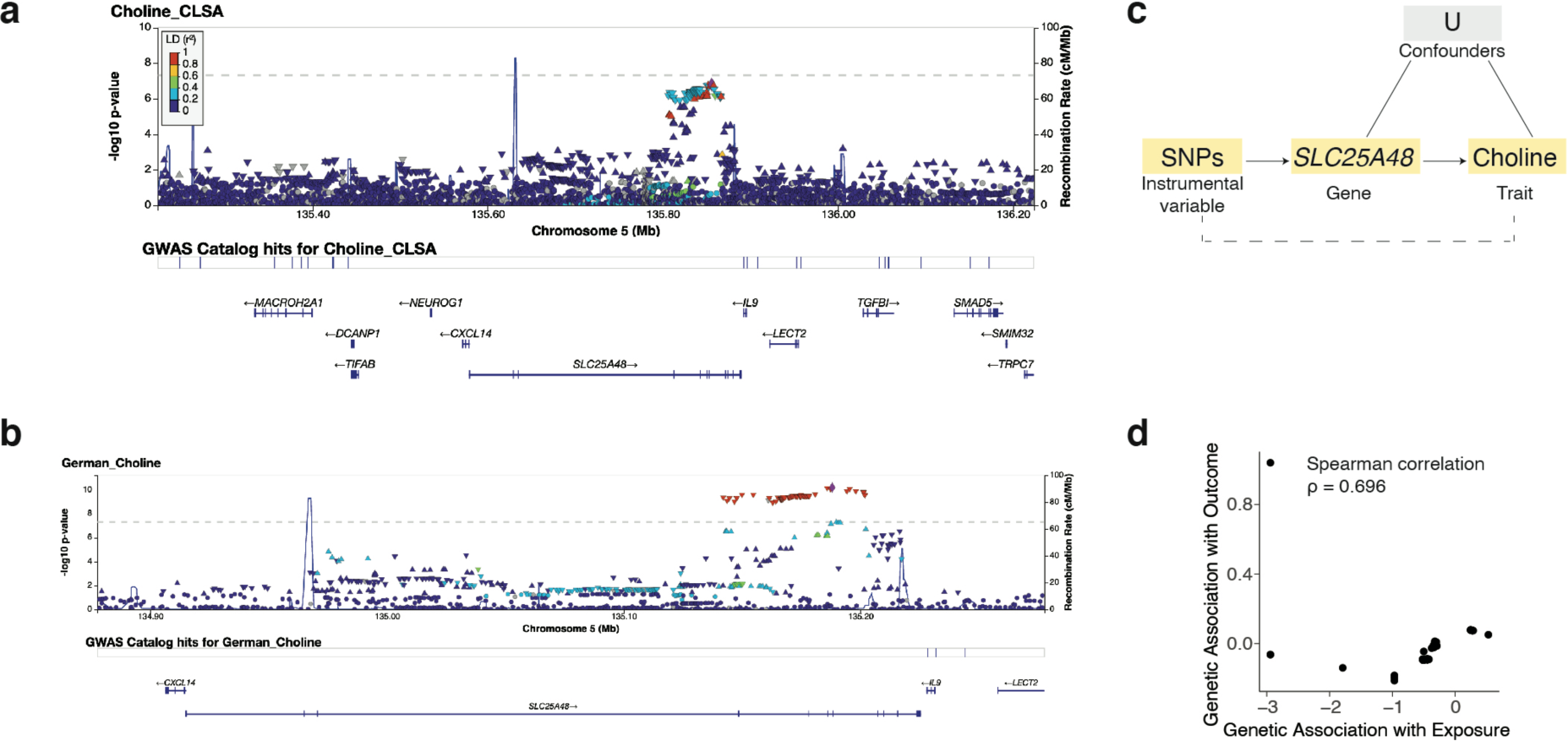
*SLC25A48* as a genetic determinant of blood choline level. a. LocusZoom plot for CLSA blood choline association signals in *SLC25A48* region. b. LocusZoom plot for GCKD blood choline association signals in *SLC25A48* region. c. Schematic for Mendelian Randomization analysis of *SLC25A48* effect on plasma choline level. d. Scatterplot showing the genetic association with *SLC25A48* viewed as exposure (x-axis) and choline viewed as outcome (y-axis). Spearman correlation ρ = 0.696.

**Extended Figure 4.**
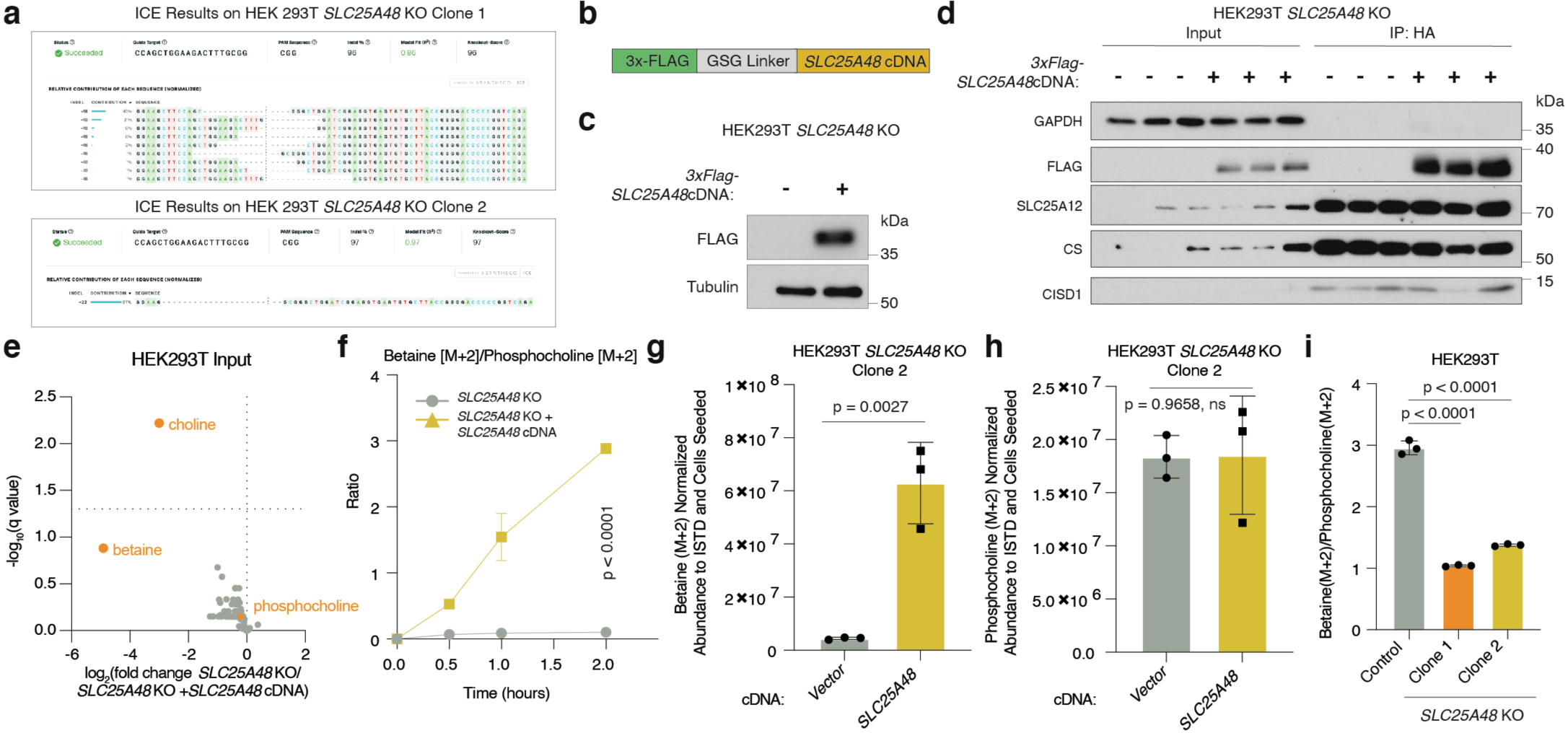
Biochemical characterization of SLC25A48 as a mediator of mitochondrial choline import. a. ICE sequencing results for the generated single clones of HEK293T *SLC25A48* knockout cells. b. Schematic of construct for *3xFLAG-SLC25A48*. c. Immunoblot of SLC25A48 (Flag) in HEK293T knockout cells expressing an empty control vector or *3xFLAG-SLC25A48* cDNA. Tubulin was used as loading control. d. Immunoblot of indicated proteins in input (whole cell) and immunopurified mitochondria from HEK293T *SLC25A48* knockout cells expressing a vector control or 3xFLAG-*SLC25A48* cDNA. e. Volcano plot with −log_10_(q-value) vs. log_2_ fold change in metabolite abundance normalized to ISTDs and protein concentration in input (whole cell) from HEK293T *SLC25A48*-knockout cells expressing a vector control or *SLC25A48* cDNA. The dotted line is the significance threshold of q-value < 0.05. Two-tailed unpaired t-tests followed by Benjamini, Krieger, and Yekutieli multiple test correction were performed. f. Betaine [M+2]/Phosphocholine [M+2] abundance ratio in HEK293T *SLC25A48*-knockout cells expressing an empty vector control or *3xFLAG-SLC25A48* cDNA after incubation with [1,2-^13^C_2_]Choline for the indicated time points. Data are mean ratio ± standard deviation. The metabolite abundances were normalized by ISTDs and cells seeded; n = 3. Statistical significance determined by two-way RM ANOVA followed by post hoc Bonferroni multiple correction. g. Barplot of betaine [M+2] abundance in HEK293T *SLC25A48*-knockout cells (Clone 2) expressing an empty vector control or *3xFLAG-SLC25A48* cDNA after incubation with [1,2-^13^C_2_]Choline for 2 hours. Data are mean ± standard deviation and normalized by ISTDs and cells seeded; n = 3. Two-tailed unpaired t-test was performed to determine statistical significance. h. Barplot of phosphocholine [M+2] abundance in HEK293T *SLC25A48*-knockout cells (Clone 2) expressing an empty vector control or *3xFLAG-SLC25A48* cDNA after incubation with [1,2-^13^C_2_]Choline for 2 hours. Data are mean ± standard deviation and normalized by ISTDs and cells seeded; n = 3. Two-tailed unpaired t-tests was performed to determine statistical significance. i. Barplot showing the ratio of metabolic abundance of betaine [M+2] to phosphocholine [M+2] in HEK293 parental and *SLC25A48* knockout cells after incubation with [1,2-^13^C_2_]Choline for 24 hours. Data are mean ratios ± standard deviation. The metabolite abundances were normalized by ISTDs and cells seeded. Two-tailed unpaired t-tests were performed to determine statistical significance.

Supplementary Table 1. Annotation of Gene-Metabolite Pairs

## Methods

### GWAS of plasma metabolites

We conducted the largest-to-date TWAS of the human metabolome. We utilized two large-scale blood metabolomic studies, leveraging GWAS summary statistics from the METSIM study^14^ (1391 metabolites in 6136 Finnish men) and the CLSA cohort^15^ (1,091 metabolite and 309 ratios, 8,299 individuals of European ancestry). Harmonization of metabolites was performed according to the study by Chen et al^15^.

### Gene-mediated regulation of metabolites

We applied S-PrediXcan^22^ independently to blood metabolites GWAS summary statistics from the METSIM and CLSA studies by using 49 JTI TWAS tissue models^23^. To calculate the replicability statistic *π*_1_^24,25^, we selected the gene-metabolite entries with q-value^25^ < 0.05 in a discovery dataset and filtered the corresponding entries in the validation dataset in a tissue-specific manner. To identify the significantly replicating gene-metabolite pairs, for each tissue model, we selected the gene-metabolite associations with q-value < 0.05 in the discovery dataset, and for the filtered subset of corresponding entries in the validation, computed q-value; the identified gene-metabolite pairs attaining q-value < 0.05 and having the same beta effect direction were chosen as significant.

### Replicability statistic and tissue sample size

The replicability statistic *π*_1_ for the gene-metabolite relationships was calculated for each tissue. The number of identified significant gene-metabolite associations in each tissue was tested for its correlation (Pearson) with the tissue sample size used in the model training^23^.

### Metabolic genes

The metabolic gene list was compiled from the Metabolic Atlas^30^, a set of transporters, and a study published by Kenny et al^6^.

### Pathogenicity scores

We utilized AlphaMissense, which builds on AlphaFold to predict the pathogenicity of variants on proteins^26^. For each gene region (boundaries defined by Ensembl GRCh38.p13), we calculated the average pathogenicity score across the missense variants. CADD scores for all possible SNVs of GRCh38/hg38 were obtained from the Washington University Server^27^. For each gene region (boundaries defined by Ensembl GRCh38.p13), we selected the maximum CADD score. The annotation of the gene as protein coding and metabolic was based on the Ensembl GRCh38.p13 annotation and the metabolic gene list, respectively.

### Genetically Determined Metabolic Networks (GDMNs)

#### Selection of associations

To build the genetically determined metabolic networks, we utilized gene-metabolite associations identified using the JTI whole blood expression model. For the nodes of the networks, we selected the metabolites that were present in at least one significant association involving metabolic genes in the discovery dataset. For the gene selection, in the discovery and validation datasets analyzed independently, we identified the list of genes scoring for the selected metabolites.

#### Building the network

For each dataset separately, based on the list of selected genes and metabolites, we built the gene-metabolite beta-effect matrix. For all the metabolite pairs, we computed Pearson correlations (link strength). To create a sparse, scale-free, and robust GDMNs with significant links, we computed the empirical p-value for each link by permutation test and then selected edges with Bonferroni-corrected p-value < 0.05.

#### Replicability assessment

To investigate the replicability of the network properties, we compared the Pearson correlations of the metabolite pairs computed from the CLSA and METSIM datasets. Additionally, we computed and correlated the node degree (number of links connected to the node) between these datasets. To study global network properties, we performed Louvain community detection and tested the community replicability between CLSA and METSIM GDMNs. The communities from the two datasets were considered overlapping if the overlap score Ω > 0.2 (Ω = (Ω_1_ ∩ Ω_2_)/min(Ω_1,_ Ω_2_), where Ω_1_ and Ω_2_ are the sets of nodes for the communities under comparison) and Bonferroni corrected Fisher’s exact test p-value < 0.001^40^.

#### Building the consensus GDMN

GDMN built on the CLSA dataset was used as the discovery dataset. To identify the significant links, we selected associations with empirical p-value < 0.05 (Bonferroni corrected) in the discovery dataset. Then, for the filtered subset of corresponding metabolite pairs in the validation dataset, Bonferroni multiple test correction of empirical p-values was performed. The entries with adjusted p-value < 0.05 were identified as significant links.

#### Scale-free property assessment

By the definition of scale-free networks, the degree distribution should follow the power-law^28^. To test whether the observed GDMN is scale-free, we computed the distribution of the node degree and showed that it follows the power-law. For comparison, we also generated 20,000 random networks with the same number of nodes and links as in the unified GDMN. The degree distribution of the random networks did not follow the power-law.

#### Error tolerance

One of the properties of a scale-free network is error tolerance. The chances of affecting the hubs are low, and disruption of a random node does not tangentially alter the global network characteristics. To assess the error tolerance, we sequentially removed 10 random nodes or hubs (from the most to less connected) and computed the mean distance between the nodes for the resulting networks. The removal of 10 random nodes was performed 500 times and the average was illustrated.

### Selection of candidate causal genes

We decided to focus on the associations involving metabolic genes due to their higher level of replicability. To narrow down to the list of the robust hits, we filtered the gene-metabolite pairs identified as significant in more than two expression models. Subsequently, to assess the power of GeneMAP to detect novel gene-metabolite relationships missed by a single-SNP analysis approach, we determined how many significant such pairs were distal (> 0.5 Mb) from significant SNPs from the CLSA GWAS.

### Annotation of selected gene-metabolite pairs

We also determined how many gene-metabolite pairs defined as “Effector” – that is, identified in CLSA through proximity to a significant SNP for the metabolite, biological relevance (i.e., participation in the biological processes of the metabolite), and colocalization with a significant cis-acting eQTL – were found with GeneMAP^15,41^. We classified the remaining significant gene-metabolite pairs as (1) “Annotated” (not identified in CLSA but very likely to be causal based on other literature), (2) “Unknown” (causal gene could not be nominated), and (3) “Miscellaneous” (associations involving uncharacterized, partially characterized compounds, or the ones with the potentially causal gene in proximity). For this classification task, we employed the annotations from CLSA^15^, Metabolic Atlas^30^, EGEA^14,16,29^, other published studies, and manual curation (Supplementary Table 1).

### Gene-metabolite pair prioritization

For each gene-metabolite pair, we selected the tissue model with the lowest p-value in the discovery dataset. We then ranked all gene-metabolite pairs by the p-value in the validation dataset. For the functional study, we focused on the gene-metabolite entries distal (> 0.5 Mb) from the significant SNPs identified in CLSA.

### Mendelian randomization

Mendelian randomization is an epidemiological approach that leverages genetic variants to detect causation (as opposed to correlation) in observational data (e.g., GWAS). To perform robust causal inference on a gene-metabolite pair, we used MR-Egger regression and the weighted median estimator^42^. We illustrate the approach by testing the SLC25A48 causal effect on choline (CLSA choline GWAS summary statistics^15^ was used). We selected the variants ±1 MB upstream and downstream according to the SLC25A48 boundary as defined by Ensembl GRCh38.p13). We then used the eQTLs from the GTEx Analysis Release V8 (dbGaP Accession phs000424.v8.p2)^43^ in this *cis*-region as genetic instruments in the Mendelian randomization analysis. To minimize weak instrument bias in Mendelian randomization, we chose the strongest eQTLs for the gene^44^.

### Loss-of-function variant analysis

We leveraged the UK Biobank whole-exome sequencing dataset in 469,787 individuals to investigate the consequences of SLC25A48 on the medical phenome. We first mapped the ICD10 code clinical data to the phecode system^45^. The phecode system has well-defined exclusion criteria to prevent contamination by cases of the control samples. We then conducted Sequence Kernel Association Test (SKAT)^46^ to test the association between LoF variants^39^ (minor allele frequency < 1%) in the gene and the disease phenotypes. In total, we tested 1,437 phenotypes in the association analysis. We used age, sex, and the first 10 principal components (PCs) derived from the genotype data (quantifying genomic ancestry) as covariates. The associations meeting FDR < 0.05 (Benjamini-Hochberg) were deemed statistically significant.

### Clinical impact of genetically determined choline level

Using PRSice 2, we trained a genetic score (PRS) of choline level and found p-value < 0.05 to be optimally predictive^47^. We then tested the resulting genetic model for choline level against the phecode-derived SLC25A48-associated disease phenotypes from the LoF analysis (p-value < 0.05) using the UK Biobank whole-genome sequencing data in 200,006 individuals. As before, we adjusted for age, sex, and the first 10 genotype-based PCs.

### Cell lines and reagents

Human cell line HEK293T was purchased from the ATCC. The cell line was confirmed to have no mycoplasma contamination and the identity was verified by STR profiling. HEK293T cells were cultured in RPMI 1640 medium (Gibco) containing 2mM glutamine, 10% FBS, 1X penicillin and streptomycin (Gibco, #15140-122).

Powdered RPMI 1640 Medium w/o L-Glutamine, L-Methionine, Choline Chloride, Folic Acid and Vitamin B12 (US Biological Life Sciences #R8999-21) was used to make choline depleted medium. After reconstitution, the following metabolites were added to match standard RPMI 1640 medium concentrations: 2mM L-glutamine (Alfa Aesar #J60573), 100.7µM L-methionine (Alfa Aesar #J61904), 2.3mM folic acid (Alfa Aesar #J62937), and 3.7nM vitamin B12 (Alfa Aesar #A14894). The medium was supplemented with 10% dialyzed fetal bovine serum (dFBS) (Gibco #26400-044) and 1X penicillin streptomycin (Gibco #15140-122).

Antibodies against HA-Tag (3724S, 1:200 for immunofluorescence), citrate synthase (14309S, 1:1,000 for Western blot) were from Cell Signaling Technology; SLC25A12 (ab200201, 1:1,000 for Western blot) was from Abcam; GAPDH (GTX627408 1:1,000) and beta-tubulin (GTX101279, 1:2,000 for Western blot) were from GeneTex; CISD1 (16006-1-AP, 1:1,000 for Western blot) was from Proteintech, Flag-Tag (F1804, 1:2,000 for Western blot, 1:200 for immunofluorescence) was from Sigma-Aldrich. Anti-mouse IgG–HRP (7076S, 1:3,000 for Western blot) and anti-rabbit IgG–HRP (7074S, 1:3,000 for Western blot) were obtained from Cell Signaling. Antibodies for immunofluorescence staining were goat anti-mouse Alexa Fluor 488 (A11029, 1:300) and goat anti-rabbit Alexa Fluor 555 (A21428, 1:300) from Invitrogen.

Other reagents: anti-HA magnetic beads (88837, Thermo Scientific Pierce); DAPI (D1306, ThermoFisher Scientific); polybrene (H9268), puromycin (P8833) (Sigma); blasticidin (ant-bl-1, Invivogen).

### Generation of knockout, knockdown and cDNA-overexpression cell lines

sgRNA for *SLC25A48* was synthesized by IDT and cloned into BsmBI linearized lentiCRISPR-v2 plasmid (Addgene, #75159) with T4 ligase (NEB #M0202). cDNA for *3xFLAG-SLC25A48* was cloned into BamHI-HF (NEB #R3136L) and EcoRI-HF (NEB #3101L) linearized pLV-EF1a-IRES-Puro (Addgene, #85132) plasmid by Gibson assembly. Mito-tag plasmids were based on pMXs-3XHA-EGFP-OMP25 (Addgene, #83356) and pMXs-3XMyc-EGFP-OMP25 (Addgene, #83355) with mCherry being replaced for the selection. The cDNAs for *3XHA-mCherry-OMP25* and *3XMyc-mCherry-OMP25* were cloned into pLV-EF1a-IRES-Blast backbone (Addgene, #85133) by using Gibson assembly. sgRNA- or cDNA-expressing vectors and lentiviral packaging vectors Delta-VPR and CMV VSV-G were transfected into HEK293T cells using XTremeGene 9 transfection reagent (Roche #636478700). The supernatant containing virus was collected 48 h after transfection and passed through a 0.45-µm filter. The transduction of target cells was performed in 6-well tissue culture plates by addition to the medium collected virus and 4 µg mL^−1^ of polybrene, followed by centrifugation at 2,200rpm for 80 min. The next day, media was changed to remove the virus. *SLC25A48* KO cells were selected by fluorescence associated cell sorting (FACS) of the GFP+ cells (lentiCRISPR-v1(GFP) and consequent single-cell cloning. For Mito-tag cells, mCherry+ cells were selected by FACS (pLV-3XHA-mCherry-OMP25, pLV-3XMYC-mCherry-OMP25). As no validated antibody is available, sg*SLC25A48*-transduced cells were tested by ICE analysis (Synthego) of genomic DNA Sanger sequencing. Cells expressing *3xFLAG-SLC25A48* were selected with puromycin. For all the mentioned constructs the matching empty vectors (without insert) were used as a control. We validated all constructs by Sanger sequencing. The nucleotide sequences are provided in the section “Nucleotide Sequences”.

### Nucleotide Sequences

Human SLC25A48_sg1 F: AAACCCGCAAAGTCTTCCAGCTGGC Human SLC25A48_sg1 R: CACCGCCAGCTGGAAGACTTTGCGG

ICE Sequencing Primer SLC25A48 F: CACCCTTATCCACCCAGAGT

ICE Sequencing Primer SLC25A48 R: GCTCTCAGAATTTAGCCCTGG

3xFLAG-SLC25A48 sg1 resistant cDNA: ATGGACTACAAAGACCATGACGGTGATTATAAAGATCATGACATTGATTACAAGGATGACGATGACAAGGGAGGTTCAGGTGGATCAGGAAGCTTCCAGCTGGAAGACTTTGCaGCcGGCTGGATCGGAGGTGCAGCCAGTGTCATCGTTGGCCACCCTCTGGACACAGTCAAGACTCGCCTGCAGGCTGGCGTTGGCTACGGAAACACCCTCAGCTGCATCCGCGTGGTGTACAGGAGGGAGAGTATGTTCGGCTTCTTCAAGGGCATGTCCTTCCCCCTCGCCAGCATTGCCGTCTACAACTCCGTGGTGTTTGGGGTCTTCAGTAACACGCAGCGGTTCCTCAGCCAGCACCGCTGCGGGGAGCCAGAGGCCAGTCCTCCCCGCACGCTGTCAGACCTGCTCCTGGCCAGCATGGTGGCCGGCGTGGTCTCTGTCGGGCTGGGAGGGCCCGTGGACCTCATCAAGATCCGGTTGCAGATGCAGACACAACCGTTTCGGGACGCCAACCTCGGTTTGAAGTCCAGGGCAGTGGCTCCTGCGGAGCAGCCAGCATACCAGGGGCCAGTGCACTGCATTACAACCATTGTGAGGAATGAGGGCCTGGCGGGGCTATACCGGGGGGCCAGTGCCATGCTGCTGAGGGATGTCCCAGGCTATTGCCTCTACTTCATCCCCTACGTGTTCCTGAGTGAGTGGATCACACCTGAGGCCTGCACAGGCCCCAGCCCCTGTGCCGTGTGGCTGGCGGGCGGCATGGCAGGAGCAATTTCTTGGGGGACAGCGACTCCTATGGATGTCGTGAAAAGTCGACTCCAAGCTGATGGGGTTTATTTAAACAAATATAAAGGTGTCCTGGACTGTATCTCCCAGAGTTACCAGAAGGAAGGTCTTAAAGTGTTTTTCAGAGGCATCACTGTGAACGCGGTGCGGGGCTTCCCCATGAGTGCGGCCATGTTCCTTGGGTACGAGCTGTCGCTGCAGGCTATCCGCGGGGACCACGCAGTGACGAGCCCATAA

*3xHA-mCherry-OMP25* cDNA: GCCACCATGTATCCCTATGACGTGCCTGATTACGCCGGCACAGGATCCTACCCCTATGATGTGCCTGACTACGCTGGCAGCGCCGGATACCCTTATGATGTGCCTGATTATGCTGGAGGGAGCGGCGTGAGCAAGGGCGAGGAGGATAACATGGCCATCATCAAGGAGTTCATGCGCTTCAAGGTGCACATGGAGGGCTCCGTGAACGGCCACGAGTTCGAGATCGAGGGCGAGGGCGAGGGCCGCCCCTACGAGGGCACCCAGACCGCCAAGCTGAAGGTGACCAAGGGTGGCCCCCTGCCCTTCGCCTGGGACATCCTGTCCCCTCAGTTCATGTACGGCTCCAAGGCCTACGTGAAGCACCCCGCCGACATCCCCGACTACTTGAAGCTGTCCTTCCCCGAGGGCTTCAAGTGGGAGCGCGTGATGAACTTCGAGGACGGCGGCGTGGTGACCGTGACCCAGGACTCCTCCCTGCAGGACGGCGAGTTCATCTACAAGGTGAAGCTGCGCGGCACCAACTTCCCCTCCGACGGCCCCGTAATGCAGAAGAAGACCATGGGCTGGGAGGCCTCCTCCGAGCGGATGTACCCCGAGGACGGCGCCCTGAAGGGCGAGATCAAGCAGAGGCTGAAGCTGAAGGACGGCGGCCACTACGACGCTGAGGTCAAGACCACCTACAAGGCCAAGAAGCCCGTGCAGCTGCCCGGCGCCTACAACGTCAACATCAAGTTGGACATCACCTCCCACAACGAGGACTACACCATCGTGGAACAGTACGAACGCGCCGAGGGCCGCCACTCCACCGGCGGCATGGACGAGCTGTACAAGCCGCGGCATCGAGGCGACGGAGAGCCTAGTGGAGTTCCTGTAGCTGTGGTGCTGCTGCCAGTGTTTGCCCTTACCCTGGTAGCAGTTTGGGCCTTCGTGAGATACCGAAAGCAGCTCTGA

*3xMYC-mCherry-OMP25* cDNA: GCCACCATGGAGCAGAAGCTGATTTCTGAGGAAGATCTGGGCACAGGATCCGAACAGAAACTGATTTCTGAGGAAGATCTGGGCAGCGCCGGAGAGCAGAAGCTGATTTCTGAAGAGGATCTGGGAGGGAGCGGCGTGAGCAAGGGCGAGGAGGATAACATGGCCATCATCAAGGAGTTCATGCGCTTCAAGGTGCACATGGAGGGCTCCGTGAACGGCCACGAGTTCGAGATCG AGGGCGAGGGCGAGGGCCGCCCCTACGAGGGCACCCAGACCGCCAAGCTGAAGGTGACCAAGGGTGGCCCCCTGCCCTTCGCCTGGGACATCCTGTCCCCTCAGTTCATGTACGGC TCCAAGGCCTACGTGAAGCACCCCGCCGACATCCCCGACTACTTGAAGCTGTCCTTCCCCGAGGGCTTCAAGTGGGAGCGCGTGATGAACTTCGAGGACGGCGGCGTGGTGACCGTGACCCAGGACTCCTCCCTGCAGGACGGCGAGTTCATCTACAAGGTGAAGCTGCGCGGCACCAACTTCCCCTCCGACGGCCCCGTAATGCAGAAGAAGACCATGGGCTGGGAGGCCTCCTCCGAGCGGATGTACCCCGAGGACGGCGCCCTGAAGGGCGAGATCAAGCAGAGGCTGAAGCTGAAGGACGGCGGCCACTACGACGCTGAGGTCAAGACCACCTACAAGGCCAAGAAGCCCGTGCAGCTGCCCGGCGCCTACAACGTCAACATCAAGTTGGACATCACCTCCCACAACGAGGACTACACCATCGTGGAACAGTACGAACGCGCCGAGGGCCGCCACTCCACCGGCGGCATGGACGAGCTGTACAAGCCGCGGCATCGAGGCGACGGAGAGCCTAGTGGAGTTCCTGTAGCTGTGGTGCTGCTGCCAGTGTTTGCCCTTACCCTGGTAGCAGTTTGGGCCTTCGTGAGATACCGAAAGCAGCTCTGA

### Immunofluorescence staining and confocal microscopy for co-localization

Sterile coverslips were coated with 50 µg/mL Poly-D-Lysine (ChemCruz #sc-136156) for 30 minutes at 37°C, and 500,000 cells per well of a 6-well plate were seeded and grown overnight. The next day, the cells were fixed in 4% PFA for 15 min, permeabilized with 1% triton X-100 in PBS, blocked in 1% bovine serum albumin (Sigma) for 30 min, and incubated at primary antibody overnight at 4°C. The following day, cells were washed in PBS twice and incubated in secondary antibodies for 30 minutes. After two washes in PBS, the coverslip was incubated in DAPI for 10 minutes, washed in PBS twice and mounted in ProLong Gold antifade mountant (Molecular Probes). For imaging, we used Nikon A1R MP multiphoton microscope with confocal modality with a Nikon Plan Apo γ 60X/1.40 oil immersion objective.

### Immunoblotting

Transmembrane buffer (10mM Tris-HCl pH 7.4, 150mM NaCl, 1mM EDTA, 1% Triton X-100, 2% SDS, 0.1% CHAPS) supplemented with protease inhibitors (EMD Millipore #535140 or Sigma-Aldrich #11836170001) was used to lyse the cells, followed by sonication. For mitochondrial immunopurification experiments, 1% Triton X-100 Buffer (50mM Tris-HCl pH 7.4, 150mM NaCl, 1mM EDTA, 1%Triton X-100) supplemented with protease inhibitors (Sigma-Aldrich #11836170001) was used for lysis. Lysates were centrifuged at maximum speed (> 14,000*g*) at 4°C, and supernatant was collected. Protein quantification was performed by using Pierce BCA Protein Assay Kit (Thermo Fisher #23227) with bovine serum albumin as a standard. 10-20% Tris-Glycine gels (Invitrogen #XP10205BOX) were used to resolve the samples. Transfer to PVDF membrane (EMD Millipore #1PVH00010) was performed with CAPS buffer (10mM CAPS, 10% ethanol). Membrane was blocked in skim milk (5% w/v) and incubated in primary antibodies at 4°C overnight. Secondary antibody incubation was in anti-mouse IgG-HRP (CST #7076) and anti-rabbit IgG-HRP (CST #7074) diluted at 1:3000 ratio in skim milk. Washes prior to secondary antibody incubation and blot development were performed with 0.1% Tween-20 Tris buffered saline. ECL chemiluminescence was used (Perkin Elmer #NEL105001EA or Cytiva #RPN2232) with autoradiography films (Thomas Scientific #1141J52) and SRX-101A Film Processor (Konica Minolta).

### Whole cell [1,2-^13^C_2_]Choline isotope tracing

For experiments with HEK293T *SLC25A48*-knockout cells expressing a control empty vector or *3xFLAG-SLC25A48* cDNA, 500,000 cells were seeded per well in 6-well plates in triplicate in choline depleted media. For experiments with HEK293T parental and *SLC25A48*-knockout cells, 250,000 cells were seeded in choline depleted media as the tracing was conducted for 24 hours. The following day, media was changed to the choline depleted RPMI with 21.5µM [1,2-^13^C_2_]Choline chloride (Cambridge Isotope Laboratories #CLM-548.01). At indicated timepoints, cells were washed twice with cold 0.9% NaCl (w/v), followed by metabolite extraction in ice cold 80:20 LC/MS grade methanol: water buffer containing ^15^N and ^13^C fully-labeled amino acid standards (Cambridge Isotope Laboratories #MSK-A2-1.2). Samples were vortexed for 10 minutes at 4°C and then centrifuged at maximum speed (> 14,000*g*) at 4°C for 10 minutes. Supernatants were dried under nitrogen and stored at −80°C. The natural isotope abundance correction was performed based on [M+2]/[M+0] ratios for the 0-minute timepoint, if available, or according to the theoretical distribution^48^.

### Polar metabolite profiling

The LC-MS analysis for the full MitoIP metabolome profiling and 24-hour whole cell tracing samples was conducted on a QExactive benchtop orbitrap mass spectrometer coupled to a Vanquish UPLC System (Thermo Fisher Scientific). Prior to injection, whole cell samples were resuspended in 60 µL of 50:50 Acetonitrile:Water, vortexed, and centrifuged at the maximum speed for 20 minutes (> 14,000*g*). External mass calibration was performed every 3 days using a standard calibration mixture. Polar extracts (5 μl) were injected onto a ZIC-pHILIC 150 x 2.1mm column (EMD Millipore) using a previously described LC-MS method^6^. The LC-MS metabolomics analysis for other whole cell tracing experiments as well as immunopurified mitochondria tracing was performed on an Orbitrap IQ-X Tribrid coupled to a Vanquish Horizon UHPLC system (Thermo Fisher Scientific). Prior to injection, whole cell samples were resuspended in 1000 µL of 50:50 Acetonitrile: Water, vortexed, and centrifuged at maximum speed for 20 minutes (> 14,000*g*). For chromatographic separation, samples were loaded on a SeQuant ZIC-pHILIC column (150 × 2.1 mm, 5 μm polymeric) and SeQuant ZIC-pHILIC Guard Kit (20 x 2.1 mm) at 40°C and eluted with the solvent system comprised mobile phase A (20mM ammonium carbonate + 0.1% ammonium hydroxide in water at pH 9.3) and mobile phase B (100% acetonitrile). The injection volume was set 5 µl, and samples were maintained at 4°C. The gradient (v/v) used was as follows: 0-22 min linear gradient from 90% to 40% B; 22-24 min: held at 40% B; 24-24.1 min: returned to 90% B; 24.1-30 min: equilibrated at 90% B at the flow rate of 150 µl/min. For the MS analysis, a HESI probe was operated in polarity switching mode, with the following source condition: spray voltage, 3.0 kV; sheath gas, 30 au; auxiliary gas, 7 au; the ion transfer tube temperature and the vaporizer temperature, 275°C. The following acquisition parameters were used for MS1 analysis: resolution, 120000; AGC target, 4e5; max injection time, 200 ms; m/z range, 55–815 Th. A pooled sample containing all biological samples was prepared and analyzed using a data-dependent acquisition method. The data-dependent MS/MS scans were acquired at a resolution of 15,000, 5e4 AGC target, auto max injection time mode, 1.6 Da isolation width, HCD activation, stepped normalized collision energy of 20, 30, 40 units, and loop count of 2. The instrument was externally calibrated using Pierce FlexMix Calibration Solution (Thermo Fisher Scientific), and the internal calibration feature (EASY-IC) was turned on. The analysis was performed using Skyline Daily v22^49^.

### Immunopurified mitochondria metabolic profiling

Mitochondrial immunopurification from HEK293T cells expressing either 3×HA–OMP25–mCherry (immunopurified mitochondria) or 3×Myc–OMP25–mCherry (background control) was conducted according to the protocol by Chen et al. with slight modifications^37^. In brief, cells were washed in a confluent 15 cm dish twice with cold saline (0.9% NaCl), collected in cold KPBS, and centrifuged at 1,000*g* for 1.5 minutes at 4 °C. The pellet was resuspended in 1 ml of KPBS, followed by homogenization with 30 strokes in a 2-mL homogenizer. Part of the homogenized sample was taken for the whole cell protein lysis and metabolic extraction. The remaining sample was incubated with 200 μl of anti-HA magnetic beads on a rotator shaker for 5 minutes at 4 °C, followed by three washes with ice-cold KPBS. Consequently, we used 10% of the bead volume for lysis with 1% Triton buffer (mitochondrial protein sample). The remaining 90% were extracted in 80% methanol containing heavy isotope labelled amino acid standards. The protein and metabolite samples were placed on a rotor for 10 minutes at 4°C, followed by 10 min centrifugation at maximum speed (> 14,000*g*) at 4°C. Metabolic samples were stored at −80°C.

### Radioactive choline uptake assay in immunopurified mitochondria

Mitochondria were immunopurified according to the above-described protocol with minor modifications. Immunopurified mitochondria bound to beads were incubated in mitochondrial uptake buffer (KPBS, 10mM HEPES, 0.5mM EGTA, 10.25uM total choline chloride, pH∼7.35) for 5 minutes at room temperature. Of the total 10.75µM choline chloride, only 100nM were radioactive ([Methyl-^3^H]-Choline Chloride; Perkin Elmer #NET109001MC), the rest was unlabeled (Choline Chloride; MP Biomedicals #67-48-1). Uptake was stopped by addition of ice-cold KPBS, followed by three washes in cold KPBS, and extracted in 80% methanol. Protein and metabolite samples were placed on a rotor for 10 minutes at 4°C, followed by 10 min centrifugation at maximum speed (> 14,000*g*) at 4°C. The metabolite extract was transferred into scintillation vials with 5 mL Insta-Gel Plus scintillation cocktail (Perkin Elmer #601339).

Radioactivity was measured with the TopCount scintillation counter (Perkin Elmer).

### [1,2-^13^C_2_]Choline tracing in immunopurified mitochondria

Mitochondrial Immunopurification was performed as outlined above. Immunopurified mitochondria were incubated in the mitochondrial uptake buffer for indicated timepoints at room temperature (KPBS, 10mM HEPES, 0.5mM EGTA, 21.5µM [1,2-^13^C_2_]Choline chloride (Cambridge Isotope Laboratories #CLM-548.01), pH∼7.35). Uptake was stopped by addition of ice-cold KPBS, followed by three washes in cold KPBS, and extracted in 80% methanol. Protein and metabolite samples were placed on a rotor for 10 minutes at 4°C, followed by 10 min centrifugation at maximum speed (> 14,000*g*) at 4°C. The metabolic samples were stored at −80°C.

### Statistical analysis and data visualization

Statistical analysis and data visualization were performed in Prism9 (GraphPad Software) or R. The details of statistical analysis for each experiment are described in the figure legends. Genetic association signal plots were performed by using LocusZoom^50^.

## Data Availability

We will provide open access to the generated results for academic use through an interactive webserver upon publication.

## Code Availability

Code used for analysis in this study will be available on GitHub upon publication.

## Acknowledgements

We thank all members of the Birsoy and Gamazon Labs for suggestions and also members of the Rockefeller University Proteomics Resource Center and the Flow Cytometry Resource Center. Some figures use modified illustrations from Servier Medical Art licensed under a Creative Commons Attribution 3.0 Unported License. A.K. is supported by a Boehringer Ingelheim Fonds PhD Fellowship. G.U. is a Damon Runyon Fellow supported by the Damon Runyon Cancer Research Foundation (DRG-2431-21). Y.L. is supported by NIH/NCI 1F99CA284249-01. T.C.K. is supported by NIH/NIDDK (F32 DK127836), the Shapiro-Silverberg Fund for the Advancement of Translational Research, and a Merck Postdoctoral Fellowship at The Rockefeller University. K.B. is supported by the NIH/NIDDK (R01 DK123323-01) and a Mark Foundation Emerging Leader Award and is a Searle and Pew-Stewart Scholar. E.R.G. is supported by NIH/NHGRI (R01HG011138), NIH/NIGMS (R01GM140287), and a Genomic Innovator Award (R35HG010718).

## Author Contributions

E.R.G., K.B. and A.K. conceived the project and wrote the manuscript. K.B. and A.K. designed experiments. A.K. developed the GeneMAP platform with input from E.R.G. A.K. performed most of the experiments with the assistance of G.U., Y.L., T.C.K. E.K conducted LC-MS analysis. E.R.G., P.L., and A.K. analyzed the phenomic datasets. E.R.G and P.L. developed the interactive portal.

